# Corelative light and electron microscopy (CLEM) image registration by blood vessel axes alignment

**DOI:** 10.1101/2025.06.23.661033

**Authors:** Y. Van den Berghe, A. Kremer, D. Verhaege, R.E. Vandenbroucke, Y. Saeys

## Abstract

Correlative light and electron microscopy (CLEM) is a multi-modal microscopy approach that contains complementary information from light (LM) and electron microscopy (EM). CLEM image alignment can be a challenging task for both the biological expert as well as for existing automated methods. This work introduces a novel method that focuses on aligning the medial axes of blood vessels in both the LM and EM modalities. Evaluation of 3D CLEM image registration problems shows that this approach obtains an accurate alignment in an intuitive and robust way. This method holds potential as a valuable tool for global and local alignment of a subset of challenging CLEM image alignment problems, both as guidance in a manual setting or in a more automated workflow.

## 1 Introduction

Microscopy is a fundamental tool for visualization in biological research Heinrich et al. [2021], Peddie et al. [2022], Kremer et al. [2015]. There has been a growing interest in multi-modal imaging techniques that exploit the assets of multiple modalities jointly Jiang et al. [2021], Teipel et al. [2015]. In the microscopy domain, light microscopy (LM) and electron microscopy (EM) can be combined such that functional information (obtained by LM) can be combined with contextual high-resolution ultrastructural visualization (obtained by EM). This multi-modal microscopy approach is called correlative light and electron microscopy (CLEM) Sjollema et al. [2012], Lippens and Jokitalo [2019], Kremer et al. [2021], Guérin et al. [2019], De Boer et al. [2015].

To optimally take advantage of this multi-modal approach, the image modalities need to be aligned using a technique called image registration Jiang et al. [2021], Zitova and Flusser [2003]. This is typically done in two steps: first a global, rough alignment is performed, followed by a local, refined alignment Avants et al. [2009]. However, CLEM image registration poses some specific challenges De Boer et al. [2015]. The two image modalities inherently show a gap in resolution, with EM resolution often being substantially higher (a factor of 10 or more) than LM resolution Kremer et al. [2021]. LM only shows the structures of interest (fluorescent labels provide the signal, while the rest of the cell/tissue remains black), while in EM all structures are visible and labeling these is not as straightforward De Boer et al. [2015]. There is an inherent symmetry in biological samples (repeating cells and structures that have a similar appearance) and an absence of a clear orientation (no ‘up’ and ‘down’ like in many other image domains). Like in medical imaging, often dense 3D volume stacks are investigated, which have a more complex processing in nature compared to 2D image registration Peddie et al. [2022]. CLEM processing is prone to local distortions due to manipulations of the sample between LM and EM acquisition (e.g., fixation, staining and dehydration for EM sample preparation and manual mounting of sample blocks (EM), pressure of a coverslip (LM) Kremer et al. [2021]). This results in linear and non-linear deformations and possible artifacts Krentzel et al. [2023]. The images could also differ with respect to the field of view (FOV), e.g. a small EM volume situated in a large LM volume. Additionally, the two image volumes could show only limited overlap.

This work proposes a method to align a subset of challenging CLEM image registration problems where an initialization is potentially difficult to find. It is based on a nicely spread structure throughout tissues, the blood vessels, and focuses on aligning the blood vessel medial axes rather than the contours. It combines a feature-based and intensity-based approach: first a set of potential global overlays is selected based on the blood vessel bifurcations present in the volume. From this set of affine transformations, a final overlay is selected based on an intensity similarity metric. Once a global overlay is obtained, the complete structures can be matched in both modalities, and further refinement can be obtained by including more bifurcation points and additional points sampled from the medial axes, using these as input for a local, non-linear transformation Bogovic et al. [2016].

The contribution of this work is to present an alternative method for determining an initial orientation by utilizing two sets of unpaired landmarks, i.e. blood vessel bifurcations found in each image modality. This approach provides valuable guidance in complex, dense 3D datasets, containing extensive symmetries and dissimilar image borders. Additionally, this global alignment algorithm can be extended towards a more automated workflow by automatically defining the bifurcation nodes through segmenting and skeletonizing the blood vessel structure. Finally, further refinement, i.e. local alignment, can be done by including more nodes from the medial axes, pairing these based on the obtained global transform and taking these as input for a non-linear transform.

This work is structured as follows: the following section outlines existing methods, Section 3 describes the datasets used to evaluate the algorithm, and describes the core concept of the algorithm. Section 4 describes the algorithm in more detail, illustrated by one of the use cases. Section 5 describes the quantitative and qualitative results. Section 6 discusses these results, noting limitations and suggesting avenues for future research.

## 2 Related work

Toolboxes with automated or semi-automated image registration methods exist, obtaining good results in many image domains. They can roughly be divided into intensity-based Klein et al. [2009], Maes et al. [2003], Chen et al. [2003] and feature-based Stamos and Leordeanu [2003], Bay et al. [2008], Calonder et al. [2010] approaches Zitova and Flusser [2003]. Scale-invariant feature detection (SIFT) is a feature-based method that searches for keys, specific points in an image, describes them, and then matches them between the two image modalities Lowe [1999]. In CLEM image registration, this method struggles with the highly multimodal content of the image: in the LM image only keys with respect to the blood vessel network are found, while in the EM image many more keys are found, with keys related to the blood vessel network not necessarily being the same because of the different appearance of structures in both modalities. Elastix is an intensity-based method that iteratively calculates a differentiable similarity metric between one image and a transformed second image and redefines a set of transformation parameters based on the gradient descent until convergence Klein et al. [2009]. It requires a good initialization for which often only an exhaustive search is possible since center-of-mass alignment methods are either not applicable or insufficient. This exhaustive search over possible translation, rotation, and scale parameters can be quite computation-intensive, especially for 3D imaging. Additionally, an optimal initialization can be missed due to the highly multi-modal aspect of the data and the presence of linear and nonlinear deformations, causing even first-choice similarity metrics for multi-modal data, such as mutual information (MI), to fail.

The highly multimodal content is the main reason why feature-based and intensity-based automated methods fail Krentzel et al. [2023]. A logical and general first step in the process of CLEM image registration is therefore to transform the problem into a pseudo-monomodal problem by extracting a structure recognizable in both image modalities, e.g. by segmenting this shared structure. Downstream processing can then be done either by these binary images or conversion to gray images (e.g. by using a Gaussian filter or calculating the distance transform). Although this simplifies the image registration problem significantly, it can still be challenging due to the size of the data, the linear and non-linear distortions, and the difference in resolution.

Another, CLEM-specific, automated approach to obtain a global transformation making use of segmented structures is to align point clouds derived from the contours of the segmented structures in both image modalities. Different algorithms exist to align these, such as the iterative closest point (ICP) algorithm or coherent point drift (CPD) algorithm Guan et al. [2018], Myronenko and Song [2010], Besl and McKay [1992]. The Autofinder module of the Icy eC-CLEM plugin is based on this approach Paul-Gilloteaux et al. [2017]. A distinction between two cases is made: a ‘similar content’ case and a ‘find smaller in bigger’ case. The ICP algorithm needs a good initialization. This is taken care of in the ‘find smaller in bigger’ case by defining regions with similar density of points to estimate the translation component and aligning the principal components of both regions to estimate the rotation component. However, their work is focused on dots, e.g. Q-dots. Although they mention the use with respect to segmentations, only a rather straightforward synthetic 2D and 3D example is given. Because of linear and non-linear deformations, and differences in resolution alignment of contours is difficult. While searching with smaller radii is an option, the case where images only partial overlap is not well covered. CLEM-Reg Krentzel et al. [2023] uses a coherent point drift (CPD) algorithm instead of a ICP algorithm to match points, extracted from automatically segmented mitochondria.

Recently also deeplearning based methods have been developed transforming the EM image to a FM-like image (e.g. DeepCLEM Seifert et al. [2020]), followed by conventional automated image registration. However, these approaches typically require large training datasets to achieve robust performance.

Due to these difficulties, the CLEM image registration process is in practice often performed manually using toolboxes that employ landmarks: pairs of points that are manually identified in both microscopy modalities Paul-Gilloteaux et al. [2017], Russell et al. [2017], Bogovic et al. [2016]. However, this process can be quite time consuming, expert-dependent, and subjective. Difficulties can be encountered on both the global (orienting one volume in the other) and the local level (defining optimal sets of paired landmarks on 2D curves or 3D surfaces in different images with different resolutions) Kremer et al. [2021].

## 3 Materials and Methods

### 3.1 CLEM datasets

Our proposed algorithm will be evaluated on three 3D blood vessel stained CLEM datasets (Figure 1). Pre-processing consists of obtaining a similar isotropic resolution for the LM and EM images. Given the lower resolution of the LM image modality, the images are adjusted to the LM resolution along the z-axis. This lower resolution is sufficient for determining a rough, global transformation.

**Figure 1.**
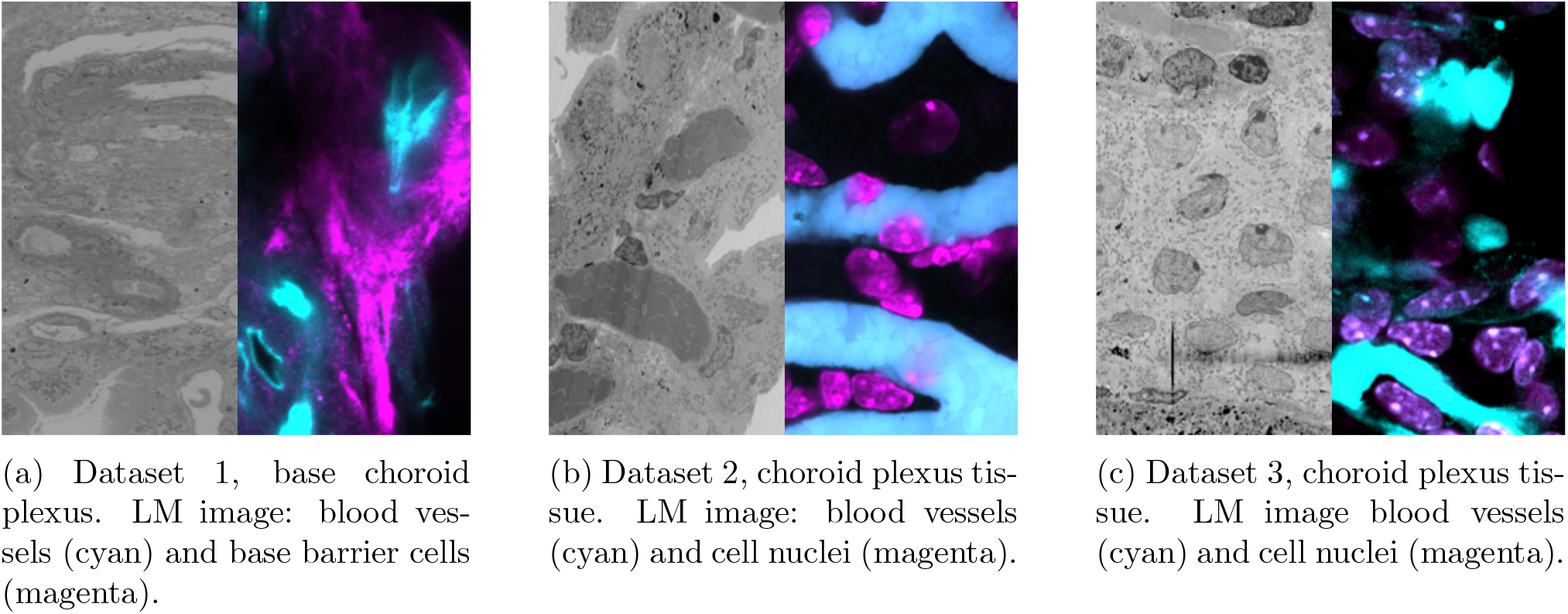
Parts of slices of the three 3D CLEM datasets containing choroid plexus tissue. In the EM image different structures can be recognized while the LM image only shows the stained structures.

The 3D datasets are generated according to the workflow described in Kremer et al. [2021]. LM data is acquired using a Zeiss LSM880 confocal microscope with Airyscan, and EM data is generated using a Zeiss Merlin SEM equipped with a Gatan 3view 2XP. The workflow involves a rotation of the sample by 90° between the acquisition of LM and EM images. As samples are mounted manually after EM sample preparation, rotation around the x and y axes is close to but not necessarily equal to zero. The z, y, and x dimensions are rather similar in EM while the z dimension is a lot smaller than the y and x dimensions in LM: the LM volumes are rather thin. All datasets were obtained from mouse choroid plexus tissue. Dataset 1 contains the region of the choroid plexus where it is attached to the ventricular walls inside the brain Verhaege et al. [2024]. This region contains specific fibroblasts (CLD11 staining, magenta), called base barrier cells, and blood vessels (IV ConA injection, cyan). Datasets 2 and 3 contain a more central part of the choroid plexus tissue in which next to the blood vessels (IV ConA injection, cyan) also the nuclei are visible (DAPI staining, magenta).

### 3.2 CLEM alignment

The datasets were aligned by an imaging expert using pairwise landmarks (BigWarp Bogovic et al. [2016]). Although primarily based on blood vessel structure, additional recognizable structures such as laser branding marks and DAPI stainings were also considered. Dataset 3 posed particular challenges as only about 50% of the LM volume and less than 10% of the EM volume showed overlapping content, making it difficult to identify similar regions in both modalities. These alignments are defined further as the ground truth (GT) alignments (third column in Figures 4 and 5 and last column in table 2). In the GT alignment, all nuclei show some overlap in the GT column since landmarks were placed on these nine nuclei. As a direct cause, they are transformed to some extent onto each other.

Initialization methods such as center-of-mass alignment, exhaustive intensity-based search, and point set density-based principal component alignment are considered inadequate for the current application due to data-specific challenges, as outlined in the Introduction. Center-of-mass alignment is often effective when the object of interest exhibits clear and symmetric boundaries, such as in structural brain MRI (magnetic resonance imaging). However, the presence of diffuse tissue structures and complex, branching vascular patterns to base image registration on renders this approach unreliable. The lack of consistent intensity-defined borders limits its applicability. Exhaustive intensity-based search methods explore a broad parameter space that includes translations, rotations, and often scaling. Despite their theoretical completeness, they often struggle with multi-modal input data (even in the pseudo-monomodal case) and the linear and non-linear deformations. The high dimensionality of the search space further exacerbates these limitations. Point set density matching followed by PC (principal component) alignment assumes that global orientation can be found in regions having similar point densities and can be corrected in a single step through rotation. This assumption breaks down in the presence of deformations and partial overlap samples. Moreover, the reliance on surface boundaries makes this method susceptible to inaccuracies in heterogeneous or irregularly shaped data. Local alignment is typically performed using intensity-based methods. However, even when initialized from an optimal or near-optimal global alignment, this step remains a challenge. While the issue of multimodality can, to some extent, be mitigated by transforming the problem into a pseudo-monomodal one, local alignment is still hindered by non-linear deformations. These introduce a highly non-convex similarity landscape with numerous local minima, often leading to suboptimal convergence. As a result, local registration may fail to capture fine-grained correspondences, particularly in regions with high anatomical variability or ambiguous intensity features.

Our proposed method focuses primarily on aligning the blood vessel backbone. Starting from two unpaired sets of blood vessel bifurcation nodes, it tries to find four paired nodes corresponding to a global (affine) transformation. Further refinement into a local alignment can then be done taking extra nodes into account. This will be explained in more detail in the next section.

## 4 MedialAxisMapping tool

Finding an affine transformation in a 3D space is equivalent to finding four paired non-linearly dependent points ((*z, y, x*) and (*z*^*′*^, *y*^*′*^, *x*^*′*^)), solving for the 12 unknowns in Equation 1.

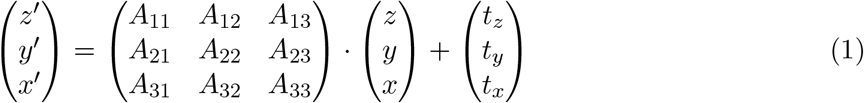

First, a set of candidate affine transformations is constructed from points representing bifurcation points in a feature-based way (steps 3 and 4 in Figure 2), after which the most optimal one is chosen in an intensity-based way (step 5 in Figure 2). Further refinement can then be obtained by including more bifurcation (and skeleton) points as input for a non-linear transformation. The algorithm will be explained in more detail below, illustrated by the datasets introduced in Section Materials and Methods, Figure 1c. An implementation of the algorithm can be found at https://github.com/yentlvdb/MedialAxisMapping. A summary of the outputs of the individual steps is given for the three datasets in Table 1.

**Table 1.** Outputs of individual steps in the global alignment algorithm for datasets 1, 2 and 3.

|  | Dataset 1 | Dataset 2 | Dataset 3 |
| --- | --- | --- | --- |
| **Global trans-form** |  |  |  |
| 1) Creation of two graphs representing the underlying blood vessel pattern | 3 EM nodes (1 EM edge) and 3 LM nodes (1 LM edge) | 8 EM nodes (5 EM edges) and 3 LM nodes (1 LM edge) | 9 EM nodes (5 EM edges) and 7 LM nodes (5 LM edges) |
| 2) Generation of all subsets of three nodes | 1 EM subset and 1 LM subset | $\binom{7}{3} = 56$ EM subsets and 1 LM subset | $\binom{9}{3} = 84$ EM subsets and $\binom{9}{3} = 35$ LM subsets |
| 3) Selection of pairs of triangles with similar edge counts | 1 pair | 24 pairs | 589 (1 edge) + 6 (2 edges) |
| 4) Tetrahedron pairs with minimal shear and scale components | 2 pairs ($thr_{sc} = 1.5$ and $thr_{sh} = 0.1$) | 2 pairs ($thr_{sc} = 1.5$ and $thr_{sh} = 0.1$) | 14 pairs ($thr_{sc} = 1.5$ and $thr_{sh} = 0.1$) |
| 5) Affine transformation matrix $A^*$ corresponding with the highest similarity metric | correct global alignment | correct global alignment | correct global alignment |
| **Local trans-form** |  |  |  |
| Added points | 6 endnodes | 5 endnodes | 4 endnodes |

**Table 2.**
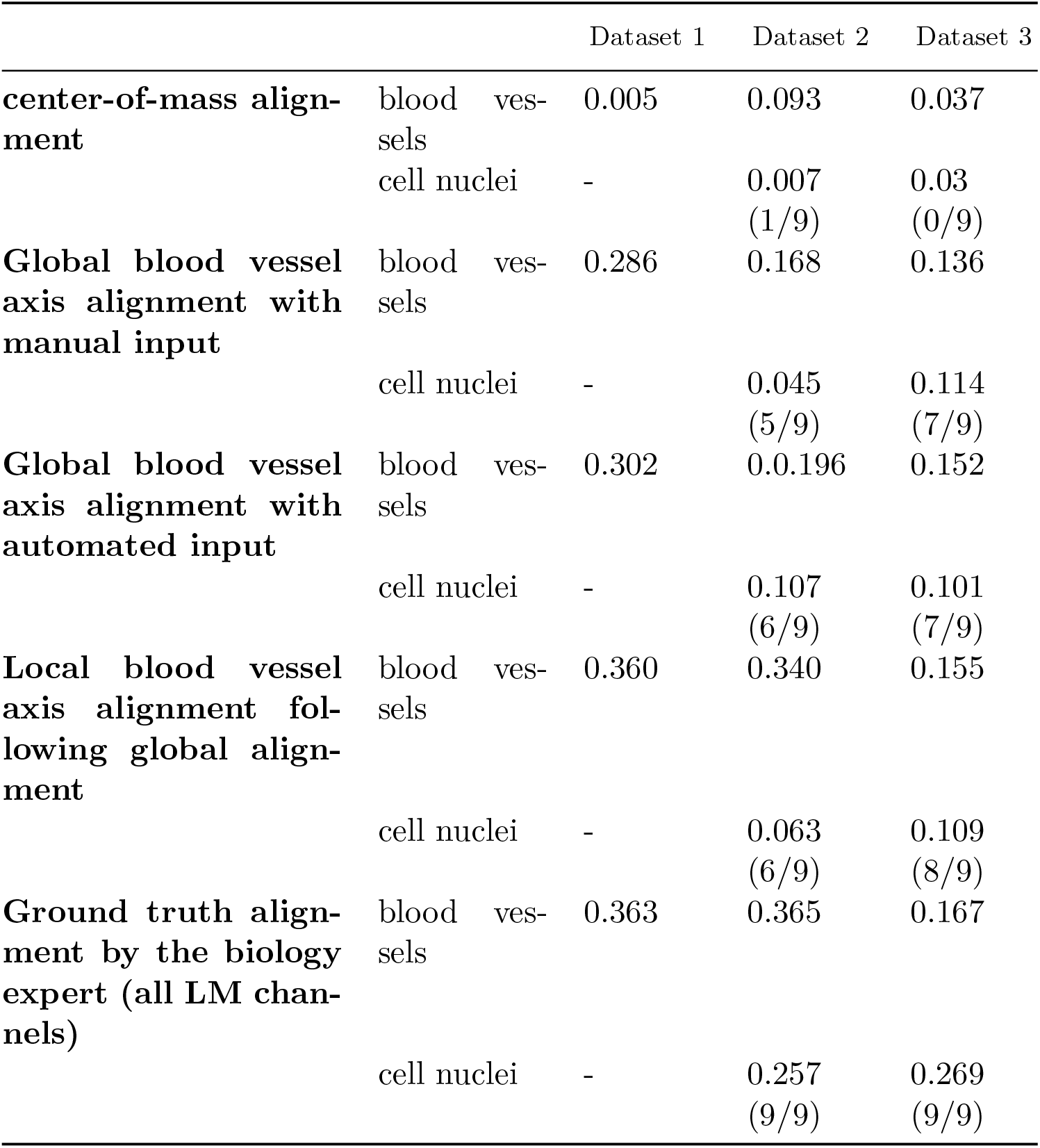
Similarity (Jaccard index) of the overlays of the three 3D CLEM datasets for both the blood vessel structure (dataset 1, 2 and 3) and the nuclei (dataset 2 and dataset 3)

**Figure 2.**
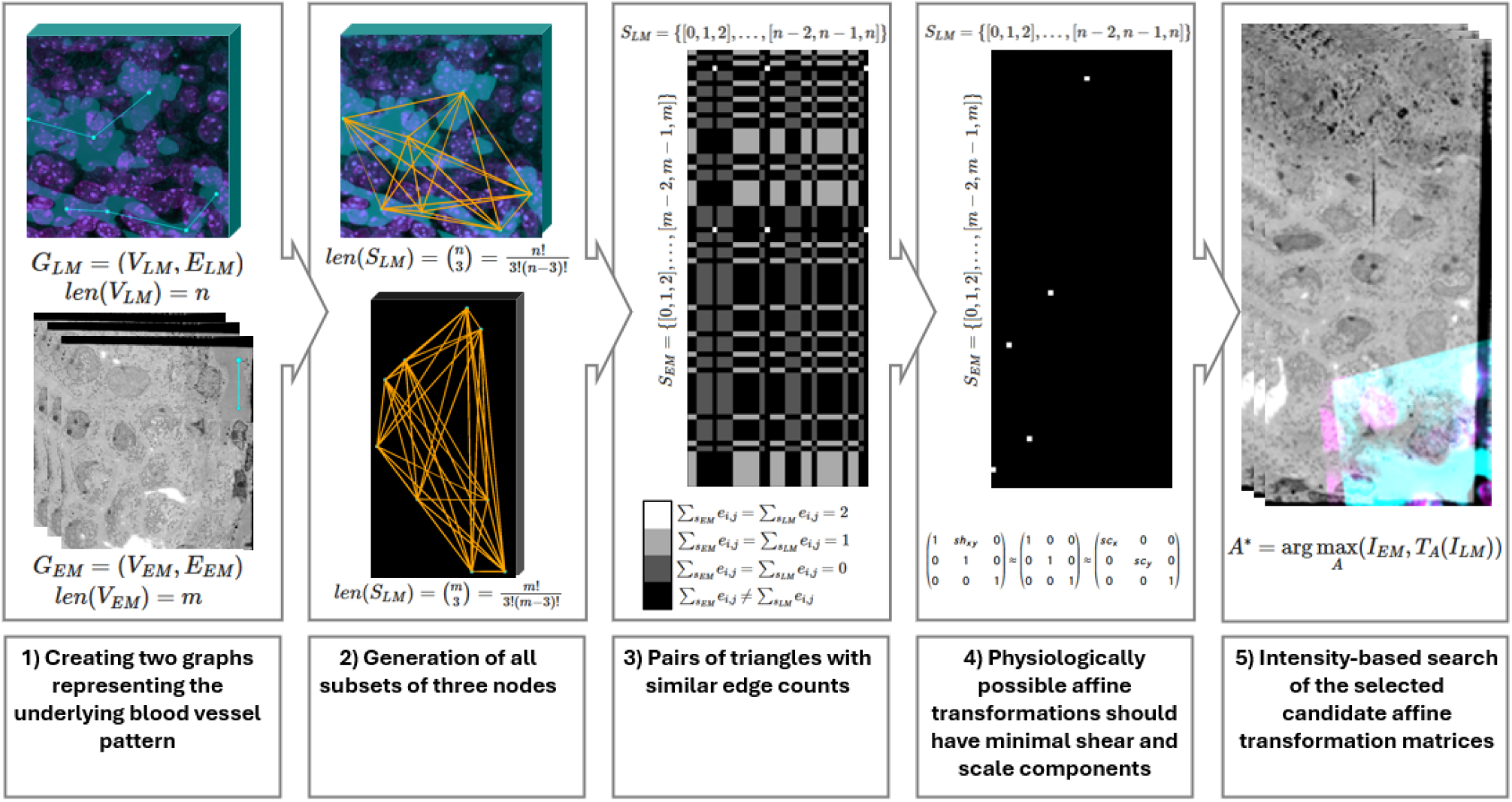
Overview of the global CLEM image alignment algorithm based on blood vessel axis alignment. 1) Two graphs representing the underlying blood vessel pattern are created. 2) Generation of all subsets of three nodes. 3) Selection of pairs of triangles with similar edge counts. 4) Determination of possible corner permutations and selection of triangle pairs with minimal shear and scale components. 5) The optimal global transformation by the affine transformation matrix *A*^*∗*^ is the transformation yielding the highest similarity metric.

### 4.1 Global transformation

The different steps of the algorithm for finding a global alignment are described in pseudo-code, see Algorithm (1), and visualized in Figure 2: 1) The process begins by creating two graphs that represent the underlying blood vessel pattern. 2) From these graphs, all possible subsets of three nodes are generated, forming all possible triangles. 3) Next, pairs of triangles with similar edge counts (0, 1, or 2 edges) are identified. 4) For each EM triangle, the possible corner permutations of the corresponding LM triangle are determined. From these, triangle pairs with minimal shear and scale components are selected as candidates for further intensity-based evaluation. 5) The transformation that yields the highest similarity metric is then chosen, and this transformation is applied to the LM image.

#### Algorithm 1 Global CLEM Image Alignment Algorithm

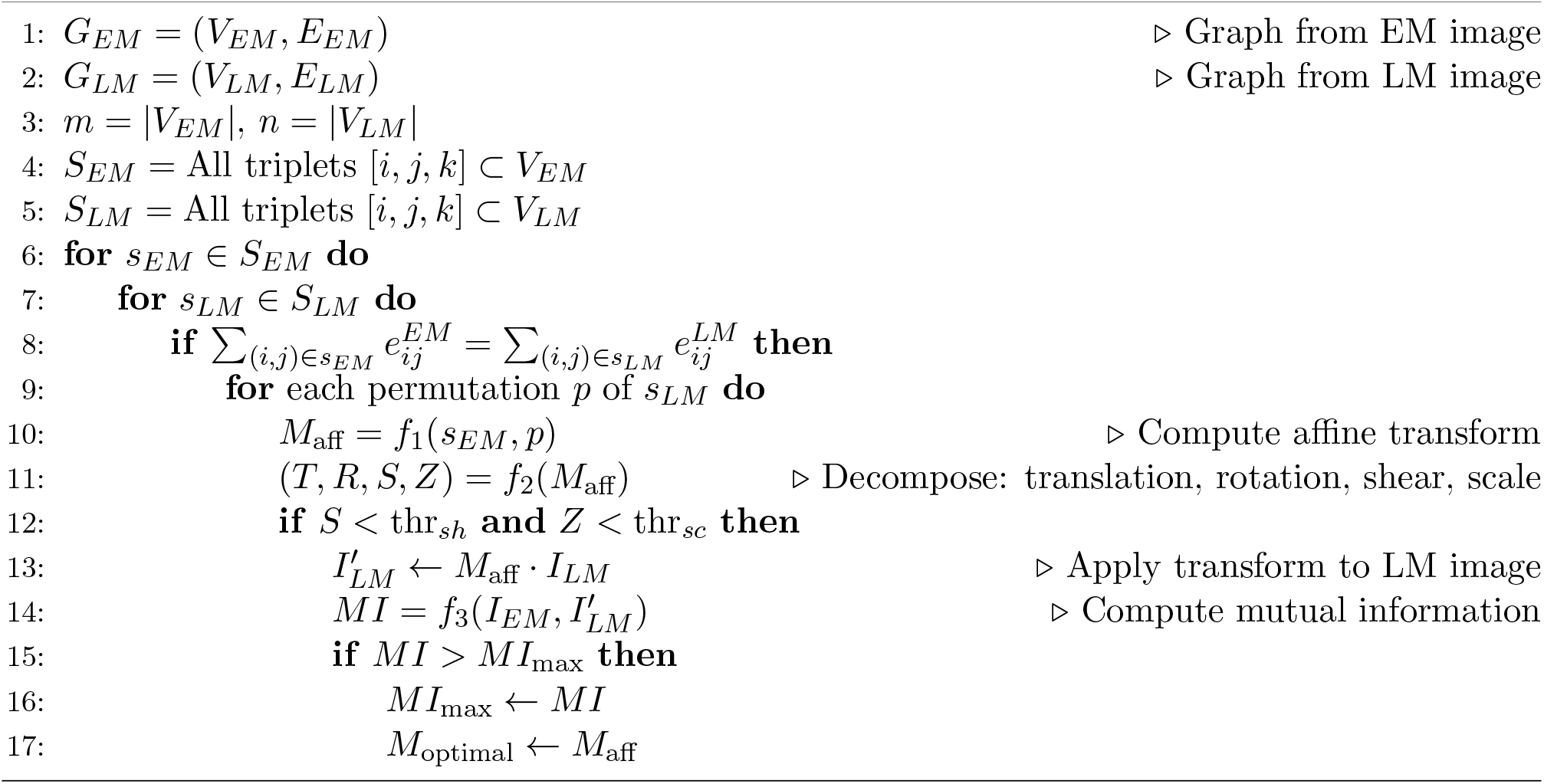

#### 1) Creating two graphs representing the underlying blood vessel pattern

Oliveira et al. [2016], Bossard et al. [2012], Drees et al. [2021]

By defining vessel bifurcations as junctions and vessel segments connecting these bifurcations as edges, two spatial global graphs are created: *G*_*EM*_ = (*V*_*EM*_, *E*_*EM*_) and *G*_*LM*_ = (*V*_*LM*_, *E*_*LM*_), with *V*_*EM*_ a set of m nodes

*{v*_*EM*,0_, *v*_*EM*,1_, …, *v*_*EM,m−*1_*}*, with *v*_*EM,i*_ = (*z*_*EM,i*_, *y*_*EM,i*_, *x*_*EM*_), a pixel in the EM image with coordinates (*z*_*i*_,*y*_*i*_, *x*_*i*_), and *E*_*EM*_ *⊆ {{v*_*i*_, *v*_*j*_*}* | *v*_*i*_, *v*_*j*_ *∈ V*_*EM*_ *}*, a set of edges, i.e. pairs of nodes.

Similarly, *V*_*LM*_ is a set of n nodes *{v*_*LM*,0_, *v*_*LM*,1_, …, *v*_*LM,n-*1_*}*, with *v*_*LM,i*_ = (*z*_*LM,i*_, *y*_*LM,i*_, *x*_*LM,i*_), a pixel in the LM image with coordinates (*z*_*i*_, *y*_*i*_, *x*_*i*_), and *E*_*LM*_ *⊆ {{v*_*i*_, *v*_*j*_*}* | *v*_*i*_, *v*_*j*_ *∈ V*_*LM*_ *}*. See Table 1 for datasets 1, 2 and 3 and the first column of Figure 2 for dataset 3 more specifically.

This step could be extended to a more automated workflow through subsequent segmentation (histogram thresholding for the LM image Yen et al. 1995] and shallow pixel classifying for the EM image Arganda-Carreras et al. [2017]) and skeletonization (Skan skeleton analysis libraryNunez-Iglesias et al. [2018]), see the first 3 columns of Figure 3. The bifurcation nodes are skeleton pixels having at least three neighbor skeleton pixels. Medial axes pixels having at most one neighbor medial axes pixel are often out-of-view nodes related to image borders and are not directly pairable as such. With regard to global alignment (column 4 of Figure 3), only the junction nodes are considered, while with regard to local alignment (column 5 of Figure 3) the end nodes are also considered. An edge is defined between (junction) nodes that are connected through part of the skeleton. However, some manual guidance might be necessary. Pruning of side branches and joining of nearby junctions should be performed adequately in order to extract an adequate graph.

**Figure 3.**
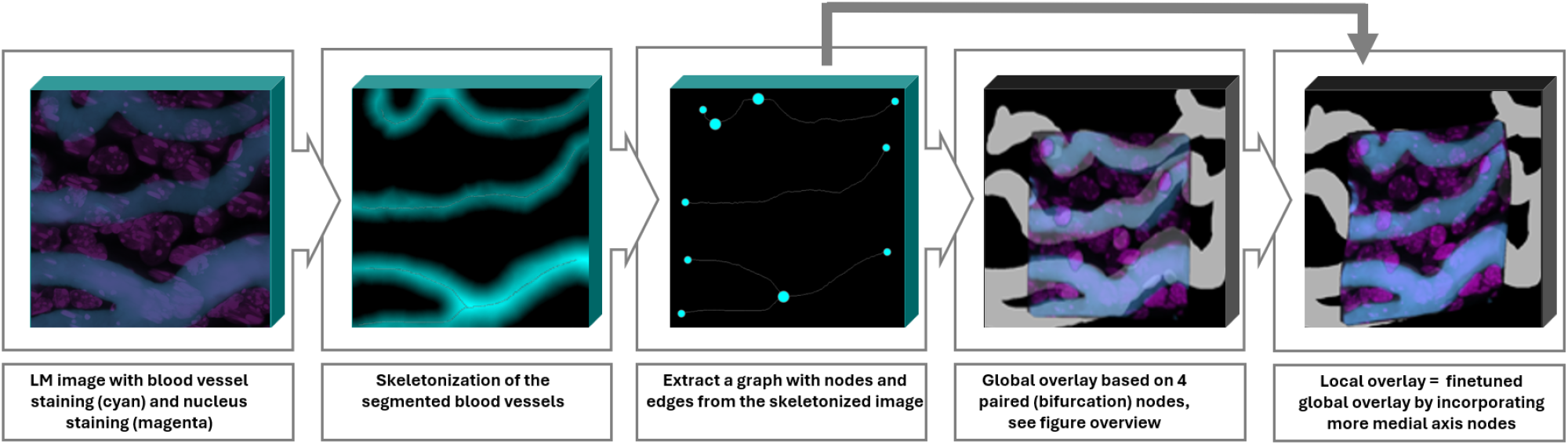
Extension towards an automated workflow. The LM image (first column) is segmented and a skeletonization is obtained through iterative thinning while keeping the original connected component intact (second column). A graph is generated for the LM image (third column). The same steps are performed on the EM image (not shown). A global transform is derived based on a subset of bifurcation nodes (fourth column, see Figure 2). Based on the remaining nodes (third column) and the global transformation (fourth column) a refined local transform can be obtained (fifth column).

### 2) Generation of all subsets of three nodes

In 3D space, finding an affine transformation matrix thus corresponds to finding four paired, non-linearly dependent points. Since in all three 3D CLEM datasets investigated here the LM modalities are rather ‘thin’ (the z dimension is significantly smaller than the x and y dimensions), the fourth point contains no extra information since it is lying in the plane described by the other three points. As a solution, three points are sufficient to estimate an affine transform, assuming no scaling or shearing exists in the direction perpendicular to the plane containing the three points. The fourth point is calculated as lying on a fixed distance on the normal, defined as 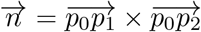of the plane, added to the center of the triangle formed by the other three nodes.

From both sets of nodes *V*_*EM*_ and *V*_*LM*_ subsets of 3 nodes are considered: *S*_*LM*_ = *{*[0, 1, 2], [0, 1, 3], [*n-* 2, *n -* 1, *n*]*}* and *S*_*EM*_ = *{*[0, 1, 2], [0, 1, 3], …, ([*m*)*-* 2, *m -* 1, *m*]*}*, w(i)th m the number of LM nodes and n the number of EM nodes. There are 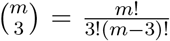 and 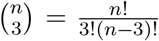 of these subsets of EM nodes and LM nodes, respectively. See Table 1 for datasets 1, 2 and 3 and the second column of Figure 2 for dataset 3 more specifically.

#### 3) Pairs of triangles with similar edge counts

These subsets of 3 points are then matched. Dataset 1 contains only one EM triangle and one LM triangle. As a result, there is thus only one triangle pair to investigate further. Dataset 2 however has multiple EM triangles, while dataset 3 has multiple EM and LM triangles. Most triangle pairs correspond to physiologically impossible deformations and should be filtered out. All triangle pairs that have a different number of edges in their subgraphs are filtered out; see the third column of Figure 2. In dataset 2, the LM triangle contains one edge. In contrast, dataset 3 includes 19, 13, and 3 LM triangles containing zero, one, or two edges, respectively. Triangles having zero or three edges result in no paired nodes and thus all six possible permutations to evaluate. In contrast, triangles having one edge or two edges result in one paired node, and thus two possible permutations to evaluate. Therefore, restricting the analysis to triangles with at least one edge substantially reduces the search space.

#### 4) Physiologically possible affine transformations should have minimal shear and scale components

Of the remaining triangle pairs, the filtered node permutations that are possible given the edge configuration, together with the calculated fourth node, correspond to a list of affine transformation matrices.

These affine transformation matrices can be decomposed into separate translation, rotation, shear and scale matrices. Since *A* is written in homogeneous coordinates such that a translation *T* can be written as a matrix multiplication, *T* can be easily extracted from *A. R* is a rotation matrix and is therefore an orthogonal matrix with the property: *R · R*^*T*^ = *I*_3_. Then a new matrix *B* = (*R · Z · S*)^*T*^ *·* (*R · Z · S*) can be defined, which can be simplified to *B* = *R*^*T*^ *· R ·* (*Z · S*)^*T*^ *·* (*Z · S*) = *I*_3_ *·* (*Z · S*)^*T*^ *·* (*Z · S*) since *R* is orthogonal, thus *B* = (*Z · S*)^*T*^ *·* (*Z · S*). *Z* is a diagonal matrix and *S* is an upper triangular matrix. The Cholesky decomposition of *B* is a decomposition of a matrix into a lower triangular matrix and its transpose: *B* = *L · L*^*T*^ with *L*^*T*^ = (*Z · S*) and *L* = (*Z · S*)^*T*^. This straightforwardly gives *Z* and *S*.

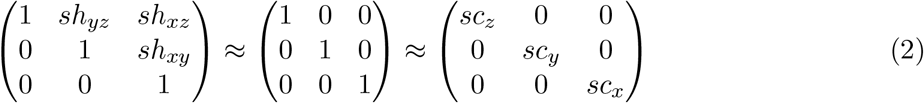

Physiologically possible transformations should have shear and scale matrices close enough to the identity matrix, thus minimal shearing and scaling, see Equation 2. All tetrahedron pairs that deviate beyond a predefined threshold should be filtered out. See Table 1 for datasets 1, 2 and 3 and the fourth column of Figure 2 for dataset 3 more specifically.

#### 5) Intensity-based search of the selected candidate affine transformation matrices

The LM image is then transformed by the remaining affine transformations, and the similarity is calculated between the EM image and the transformed LM image. The similarity metric of choice is mutual information Maes et al. [2003], Chen et al. [2003], calculated from the original images or from the Gaussian blurs of the segmented images if these are available. Mutual information between two images describes how much information of one image is contained in the other image, or thus how the pixel-wise intensity values in one image can predict the pixel-wise intensity values in the other image. The higher this value, the better the two images align. Normalized mutual information (NMI) is calculated as follows:

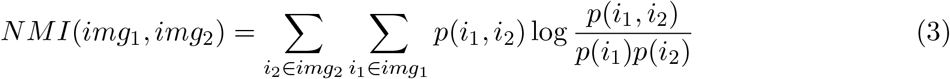

with *p*(*i*_1_, *i*_2_) the joint histogram and *p*(*i*_1_) and *p*(*i*_2_) the individual histograms. The NMI is thus high when the signal in the joint histogram is highly concentrated in a small number of bins. Incorporation of individual histograms normalizes mutual information to values between 1 (low similarity) and 2 (high similarity).

### 4.2 Local transformation: final refinement by including more points

Once a global transformation is obtained, the other bifurcation nodes can be paired based on a distance function (such as the Euclidean distance), and the graph representation. Other bifurcation nodes, if present, are paired and added to the set of paired nodes. Pairwise points are extracted from the medial axes in regions where the blood vessel structures aligned less well. The paired bifurcation nodes, together with paired medial axes nodes in the automated workflow, are then given as input to a local alignment algorithm that uses a thin plate spline approach to warp the image based on point correspondences Bogovic et al. [2016] (see Table 1 and Figure 3 for dataset 2 more specifically).

## 5 Results

The alignment method proposed in this study focuses on aligning the blood vessel network, a structure visible in both image modalities. In a first step (global alignment), an affine transformation matrix is constructed from paired bifurcation nodes. The bifurcation nodes are either manually selected or automatically derived. In a second step (local alignment), the remaining bifurcation nodes and additional skeleton nodes, if a skeletonization of the blood vessel network is available, are matched. The alignment of the EM and LM images of dataset 1 resulted in the discovery of a new barrier located in the region where the choroid plexus is attached to the brain parenchyma. Blood vessel-based alignment made it possible to recognize LM-stained fibroblasts in the EM image, revealing their clustering and the resulting sealing of the choroid plexus stroma of the brain parenchym Verhaege et al. [2024].

There are three main points of interest that will be addressed in the subsections below. First, we examine whether the proposed algorithm is effective in identifying a good global transformation when provided with manually selected, unpaired input. Second, we evaluate the algorithm’s performance in determining a suitable global transformation within an automated workflow. Third, we assess the extent to which alignment improves when additional nodes from the blood vessel medial axes are incorporated.

The alignments will be evaluated both quantitatively and qualitatively, ideally on other structures in addition to the structure on which registration is based (the blood vessel structure). However, to evaluate the accuracy of the global alignment, the evaluation of only the blood vessel structure is sufficient to determine whether the correct region is recognized. For a qualitative evaluation, see Figures 4 (2D slice of the volume) and 5 (3D view). Because the EM volume is dense, only segmented blood vessels are visible. For a quantitative evaluation, see Table 2. The Jaccard Index (JI) is used, formulated as the intersection of two binary images *img*_1_ and *img*_2_ over the union of these two images, with *TP* the number of true positives, thus where a pixel or voxel in the overlayed image has a one in both modalities, and *FN* and *FP* the number of false negatives and false positives where a pixel or voxel in the overlayed image has a one in one image modality but a zero in the other modality:

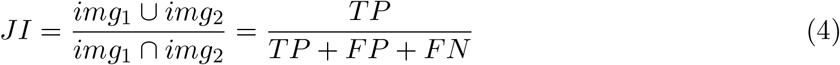

**Figure 4.**
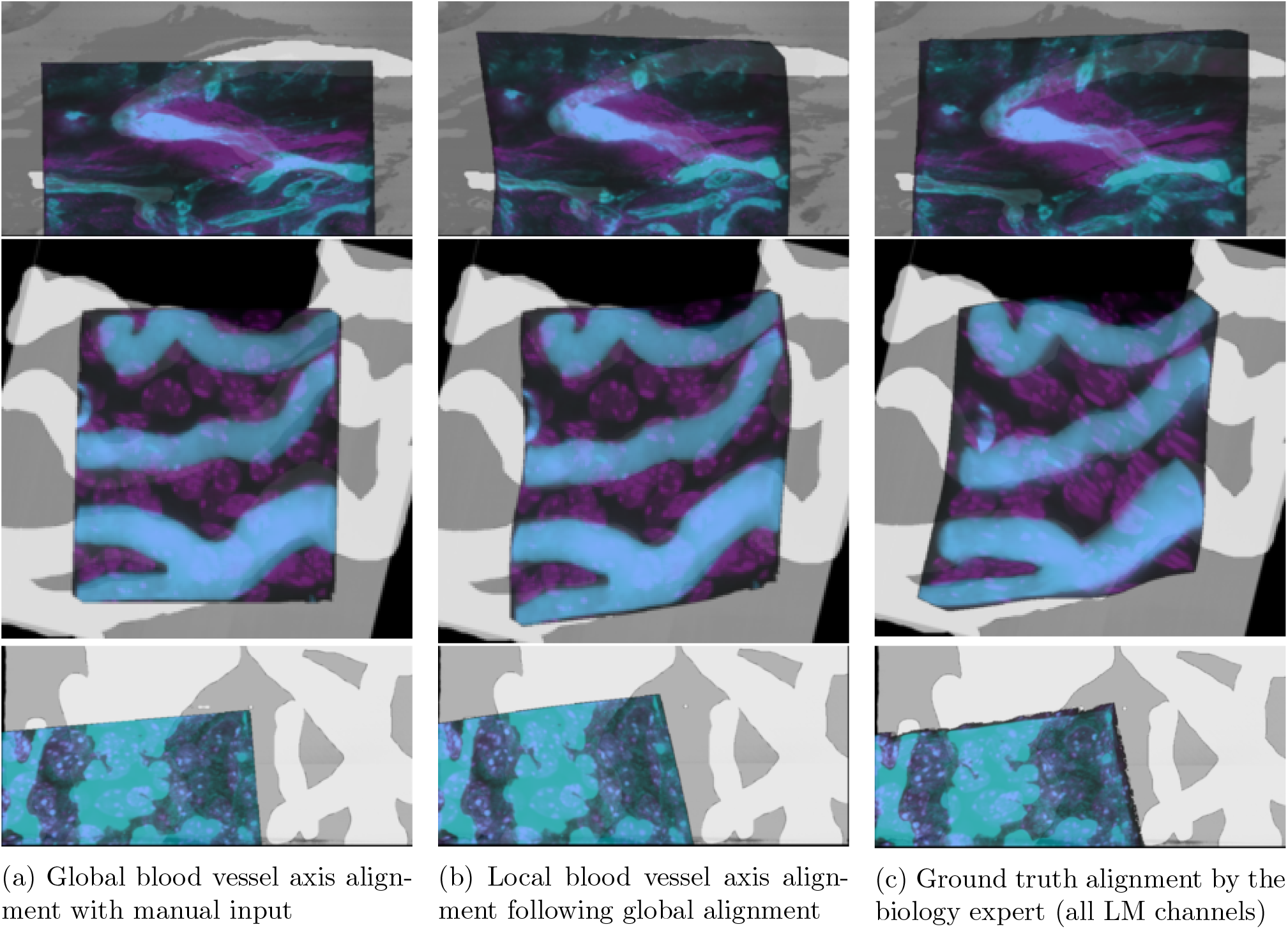
The 3D overlays are shown for the three 3D CLEM datasets: dataset 1 (first row): with LM: blood vessel (cyan) and fibroblast (magenta) staining, dataset 2 (second row): with LM: blood vessel (cyan) and nucleus (magenta) staining and dataset 3 (third row): with LM: blood vessel (cyan) and nucleus (magenta) staining. Of the EM volume, only the blood vessel structure is visible through segmenting the images.

### 5.1 Global alignment with manual input

The first column of Figures 4 and 5 shows that the corresponding regions are found using the blood vessel axis alignment method with manual input in the three 3D CLEM alignment problems. This is also reflected in Table 2 where the Jaccard indices of the blood vessel overlays are comparable to the ground truth alignment and a subset of nuclei overlap. In dataset 1 (Table 1 and the first row of Figures 4 and 5) there are not many bifurcation nodes, and thus not many combinations to consider. An optimal global alignment is rather straightforwardly selected from the two candidate affine transformation matrices. The EM volume in dataset 2 (Table 1 and the second row of Figures 4 and 5) contains more bifurcations, increasing the number of potential triangle pairs. After selecting possible triangle pairs, again only two possible overlays are selected, from which an optimal global alignment is selected based on the MI of the original images. In dataset 3 (Table 1 and the third row of Figures 4 and 5)), both the EM and LM volumes contain more bifurcations, and the number of potential triangle pairs increases considerably. To reduce this number, only triangle pairs having minimum one edge were considered, resulting in 14 pairs to evaluate. From these selected affine transformation matrices, a correct global alignment was again found.

**Figure 5.**
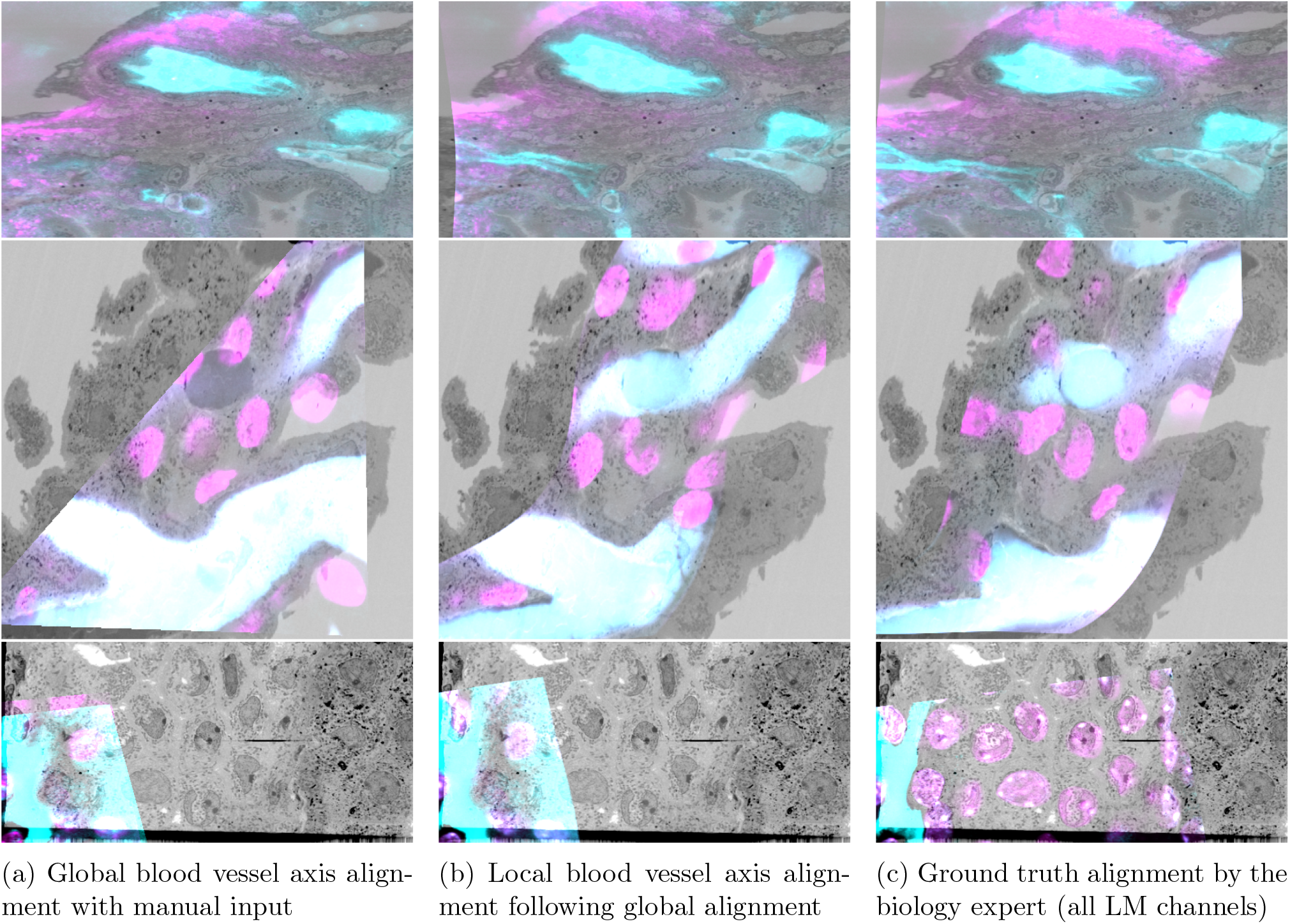
Of the alignments of the three 3D volumes a crop of an EM slice with according transformed LM image is shown: dataset 1 (first row): with LM: blood vessel (cyan) and fibroblast (magenta) staining, dataset 2 (second row): with LM: blood vessel (cyan) and nucleus (magenta) staining and dataset 3 (third row): with LM: blood vessel (cyan) and nucleus (magenta) staining.

### 5.2 Global alignment with automated input

The global alignment with automated input, thus where the bifurcation nodes were obtained in an automated way, is similar to the one obtained with manual input, see the first column of Figure 4, and the same workflow was followed (Table 1). Similar or slightly better similarity scores were obtained compared to the algorithm with manual input (Table 2).

Table 3 compares the proposed algorithm with an exhaustive approach with respect to the number of times the similarity of the aligned images is calculated, which is the most computationally intensive step due to the high number of repetitions and the size of the images. For the exhaustive method, steps of 10 voxels were taken with partial overlap up to half of the smallest image. Rotation around the z axis is done with steps of 10°, and rotation around the x and y axes is done for -10°, 0°, and +10°, since rotation around these axes is assumed to be small due to the workflow followed in Kremer et al. [2021], but can be non-zero. The number of steps in the z, y, and x directions, together with the number of rotations, is listed in the second column of Table 3, the resulting number of times the similarity of the overlay is calculated in the third column. In the proposed algorithm, while the number of possible triangle pairs can increase fast (fourth column), these numbers decrease in a first step by only considering pairs having similar edge counts (fifth column) and in a second step by only considering triangle pairs having realistic deformations, i.e. realistically low shears and scales (sixth column). The calculations up to this step involve, compared with the image matrix size, rather small (4 by 4) matrices, and are thus less computationally intensive. Of these resulting affine transformation matrices, the similarity is calculated of the overlays.

**Table 3.**
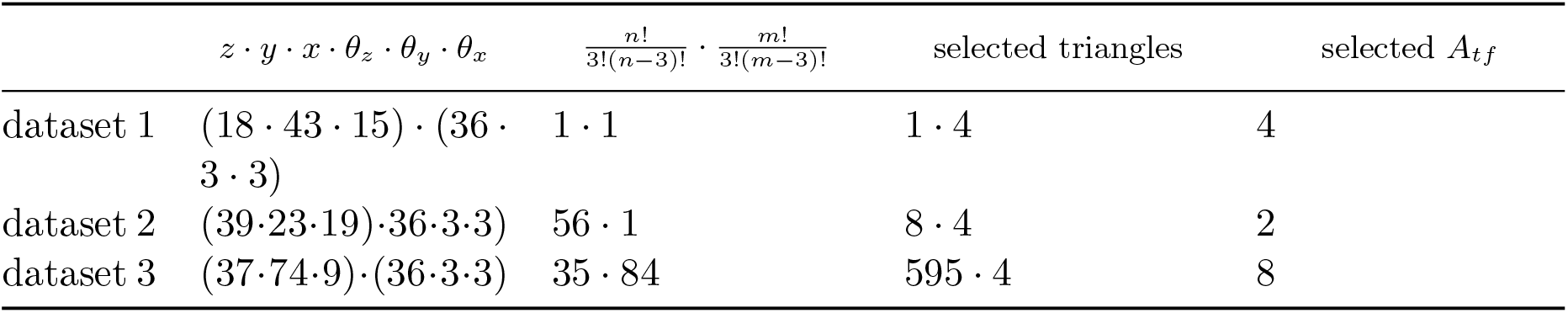
Number of times a similarity metric is calculated: 1) in a basic exhaustive way (second column), calculated for steps of 10 pixels in z, y and x directions (*z · y · x*), and steps of 10 for rotation around z, y and x axis (*θ*_*z*_ *· θ*_*y*_ *· θ*_*x*_), 2). The third column lists the number of triangle pairs investigated. The fourth column the number of triangles meeting the edge conditions times the number of node permutations, and finally sixth column, the final number of affine transformation matrices further investigated in an intensity-based way. The third and sixth column are more computationally intensive because of calculations involving the large 3D images, the fourth and fifth column are less computationally intensive because of calculations involving the small 3D affine transformation matrices.

| | $z \cdot y \cdot x \cdot \theta_z \cdot \theta_y \cdot \theta_x$ | $\frac{n!}{3!(n-3)!} \cdot \frac{m!}{3!(m-3)!}$ | selected triangles | selected $A_{tf}$ |
| --- | --- | --- | --- | --- |
| dataset 1 | $(18 \cdot 43 \cdot 15) \cdot (36 \cdot 3 \cdot 3)$ | $1 \cdot 1$ | $1 \cdot 4$ | 4 |
| dataset 2 | $(39 \cdot 23 \cdot 19) \cdot 36 \cdot 3 \cdot 3$ | $56 \cdot 1$ | $8 \cdot 4$ | 2 |
| dataset 3 | $(37 \cdot 74 \cdot 9) \cdot (36 \cdot 3 \cdot 3)$ | $35 \cdot 84$ | $595 \cdot 4$ | 8 |

### 5.3 Local alignment

Figures 4 and 5 show that the alignment improves when more paired nodes (Table 1) are added compared to global alignment. The 3D views in Figure 4 show that if the ends are misaligned in the global automated (and manual) overlays, they do align more in the local overlay. This is most visible in dataset 2 and the upper structure in dataset 3. This is reflected in Table 2 where the JI of the blood vessel segmentations is higher than that of the global alignments. The JI of the cell nuclei generally follows the same trend.

## 6 Discussion

In this study, the problem of CLEM image alignment was investigated. This can be a challenging problem for both the biological expert to perform manually as well as existing automated methods, due to the highly multi-modal aspect, linear and non-linear deformations and the possibility of only partial overlapping volumes. In this study a method is proposed that focuses on aligning the blood vessel backbone, a structure randomly distributed throughout the images. The results show that it succeeds in finding a correct global transformation both with manual and automated input, and that the local transformation further finetunes the global alignment.

Aligning image modalities by aligning blood vessel backbones is an intuitive and robust approach. It can help guide the biological expert in finding the corresponding regions when the initial orientation is not immediately clear. Pairing two sets of unpaired bifurcation nodes is often a more simple task than pairing two sets of more general landmarks, since the assumption can be made that the number of bifurcations in an image will be smaller than the number of possible landmark locations. Additionally, aligning the medial axes should be more correct compared to aligning parts of the contours because of the large gap in resolution between the two modalities. These findings are reflected in the results, where only three points in a complex 3D alignment problem already give a quite good approximation of the overlays obtained by the higher number of landmarks landmarks used in dataset 1, 21 landmarks used in dataset 2, 10 landmarks used in dataset 3). This method thus could be interesting in guiding the biological expert in a challenging image registration problem by selecting a subset of potentially adequate overlays from which the expert can select the most optimal one.

Similarly to the manual input workflow, it seems intuitively more accurate to align the medial axes instead of the contours for the automated workflow. The method proposed here also includes the case of partial overlap, which is not fully covered by existing automated methods. The eC-CLEM auto-finder method mainly covers the cases of similar content and the cases where a smaller volume fits in a larger volume (with some room for partial overlap because of the half-radius approach) Paul-Gilloteaux et al. [2017]. Exhaustive intensity-based methods can incorporate partial overlays, but the number of possible parameter combinations increases fast, making it easier to overlook an optimal local maximum. The method combines a feature-based approach with an intensity-based approach: a selection of potential global overlays is made based on landmarks, i.e. bifurcation nodes, and from this considerably smaller number of selected candidates (compared to an exhaustive approach), a final global transform is selected based on the similarity between the voxel-wise intensities.

There are some limitations present. In the datasets presented here, the number of bifurcations is rather low; between 3 and 9. It would be interesting to investigate this approach for datasets having larger numbers of bifurcations as well. While the number of possible triangles increases rapidly (*n ∗* (*n -* 1) *∗* (*n -* 2) *∼ n*^3^), the number of possible combinations in an exhaustive search also increases rapidly, in general *n << x, y, z*. The number of bifurcations is thus substantially smaller than the size of the sample. Additionally, while the exhaustive search acts on the full, large images, the algorithm first selects a subset of potential overlays based on the much smaller, 4 by 4 (3D) affine transformation matrices. Local alignment of the bifurcation nodes of one dataset with around 20 bifurcation nodes is promising, but more datasets are necessary to test this hypothesis and are being generated at the moment. Additional filter methods to select a subset of potential overlays could be investigated. Triangle pairing could be done dependent on the size of the simplexes. Other triangulation methods could be investigated such as a Delaunay triangulation of space assuming a minimal amount of similarity is present if linear and nonlinear deformations are not too profound. If less than 3 or 4 bifurcations (with a minimum of one) are present, additional points could be sampled from the blood vessel segments leaving the bifurcation. Another limitation (for the automated input approach) is the inherent dependence on segmentations, which can be a challenging and computationally intensive task. However, extracting similar structures in both images, in general in the form of deriving segmentations of those structures, is a general first step for most existing automated methods, given the highly multimodal aspect of CLEM data. Finally, in addition to applying this method on blood vessel stained data with a higher number of bifurcations, it would be interesting to test this approach with other network-like structures such as neural networks.

## 7 Conclusion

CLEM image registration can be challenging both for the biological expert and for existing automated methods. The expert may have difficulty finding similar regions in both modalities and, once found, in defining precise landmarks on 3D surfaces. Automated methods struggle with the highly multi-modal aspect, resolution differences, and large 3D volumes. The method proposed here combines a feature-based approach to select candidate overlays and an intensity-based approach to identify an optimal global overlay from the set of selected candidates. It does so by focusing on aligning the blood vessel network, a ubiquitous structure nicely distributed throughout the tissues. This method can guide the expert in finding the corresponding regions and is expandable to a more automated setting by automatically obtaining bifurcation points from skeletonized segmented structures. The global transform can then be further fine-tuned by pairing other bifurcation nodes and sampling additional paired nodes from the medial axes. This method appears to be helpful in a subset of challenging CLEM image problems and could be an interesting approach to align other challenging datasets, both as a guide in manual alignment and as an alternative approach to an exhaustive intensity-based global alignment.

## Acknowledgements

The FLanders Artificial Intelligance Research program (FAIR).

## Conflict of Interest

The authors declare no conflict of interest.

## Notes

https://github.com/yentlvdb/MedialAxisMapping

### Competing Interest Statement

The authors have declared no competing interest.

### Funding

- Flanders aritificial intelligence research program

